# NMRdock: Lightweight and Modular NMR Processing

**DOI:** 10.1101/679688

**Authors:** Kyle W. East, Andrew Leith, Ashok Ragavendran, Frank Delaglio, George P. Lisi

**Author notes:** **Corresponding Author:** George P. Lisi.

## Abstract

NMR is a widely employed tool in chemistry, biology, and physics for the study of molecular structure and dynamics. Advances in computation have produced scores of software programs necessary for the processing and analysis of NMR data. However, the production of NMR software has been largely overseen by academic labs, each with their own preferred OS, environment, and dependencies. This lack of broader standardization and the complexity of installing and maintaining NMR-related software creates a barrier of entry into the field. To further complicate matters, as computation evolves, many aging software packages become deprecated. To reduce the barrier for newcomers and to prevent deprecation of aging software, we have created the NMRdock container. NMRdock utilizes containerization to package NMR processing and analysis programs into a single, easy-to-install Docker image that can be run on any modern OS. The current image contains two bedrock NMR data processing programs (NMRPipe and NMRFAM Sparky). However, future development of NMRdock aims to add modules for additional analysis programs to build a library of tools in a standardized and easy-to-implement manner. NMRdock is open source and free to download at https://compbiocore.github.io/nmrdock/.

---

Nuclear magnetic resonance (NMR) has been one of the most valuable tools at the interface of chemistry, biology, and physics for the past 60 years. NMR can probe both the structure and dynamics of a target molecule with atomic resolution and has become ubiquitous in structural biology,^*1–4*^ synthetic^*5, 6*^ and medicinal chemistry,^*7–9*^ and materials sciences.^*10, 11*^ This utility is facilitated by a rapid improvement in technology (*i.e.* cryoprobes,^*12, 13*^ high field magnets,^*14*^ MAS/high-speed rotors^*15, 16*^) and pulse sequence design (*i.e.* TROSY,^*17, 18*^ CRINEPT,^*19, 20*^ NUS,^*21*^ Ultrafast^*22*^) that has created avenues for studies of increasingly large and complex systems. An equally important advance in the NMR field has been the improvement in computational power that allows for faster and more consistent processing and analysis of NMR data.^*23*^

Spectroscopists have thrived in this environment, designing and implementing scores of software packages catalogued by the BioMagResBank.^*24*^ Many of these programs make up the bedrock of modern NMR spectroscopy,^*25, 26*^ allowing for the processing of Fourier and nonuniformly sampled data,^*27*^ robustly fitting NMR spin relaxation parameters to extract information on chemical dynamics,^*28–30*^ and adapting NMR distance constraints to elucidate biomolecular structures.^*31*^ However, as the demand for NMR spectroscopy in biological settings grows, three significant issues have arisen: 1) the 100+ unique software packages reported in the literature have their own system-dependent environments, paths, and dependencies, complicating installation; 2) software development and utilization has been a laboratory-dependent process, with many research groups supporting different operating systems and relying on different software packages and 3) planned updates to modern operating systems are removing or altering libraries that are required by many older packages, forcing the deprecation of these programs. All three of these computational problems place large barriers for entry into the field.

Several projects attempting to reduce this barrier are underway, including those focused on automated data processing programs with helpful GUIs and powerful algorithms (*i.e.* MestreNova, CRAFT),^*32*^ transitions to system-independent languages (*i.e.* Python-based software),^*28*^ or the aggregation of software packages into a single source (*i.e.* NMRbox).^*33*^ Each of these avenues has made NMR more accessible to the greater scientific community, but there are also disadvantages to each solution. For example, advanced GUIs are not always open-source and require expensive licenses, while system-independent languages have reduced installation complexity of NMR software but still require additional integration and knowledge of proper dependencies. At the moment, a large suite of NMR software is available in NMRBox,^*33*^ a highly versatile and powerful virtual machine (VM) that integrates available software into a single unit. We sought to build upon this approach by creating a lightweight, standardized and modular containerized system in order to 1) make it easy to install and use for new spectroscopists, 2) create a long-term solution for deprecating software, and 3) make it an accessible educational tool for NMR data processing and analysis in classroom and workshop settings. In this work, we highlight the potential of containerization for implementing NMR software with no dependence on the configuration of the host.

## Why Containerization?

Our goal was to create a lightweight, easy-to-use, and adaptable tool for maintaining NMR software that can 1) run on any modern operating system (OSX, Windows, and Linux), 2) support 32-bit and deprecating software packages, and 3) maintain a modular structure to allow for future development. To meet all of these requirements, we turned to containerization.

The current standard in the field relies on virtual machines that abstract the hardware and run a guest operating system on this virtual hardware. This provides unparalleled versatility and security, but comes at the cost of massive computational overhead required to create the virtual hardware, a burden that is eased with the lightweight containerization paradigm. Unlike virtualization, containers package applications into an optimized environment, holding the program and necessary runtime libraries, but utilize the host operating system through the container engine (Figure 2). This structure is more lightweight and adaptable than a VM, and is becoming more widely employed by server-based companies like Google and Amazon.^*34*^

**Figure 1.**
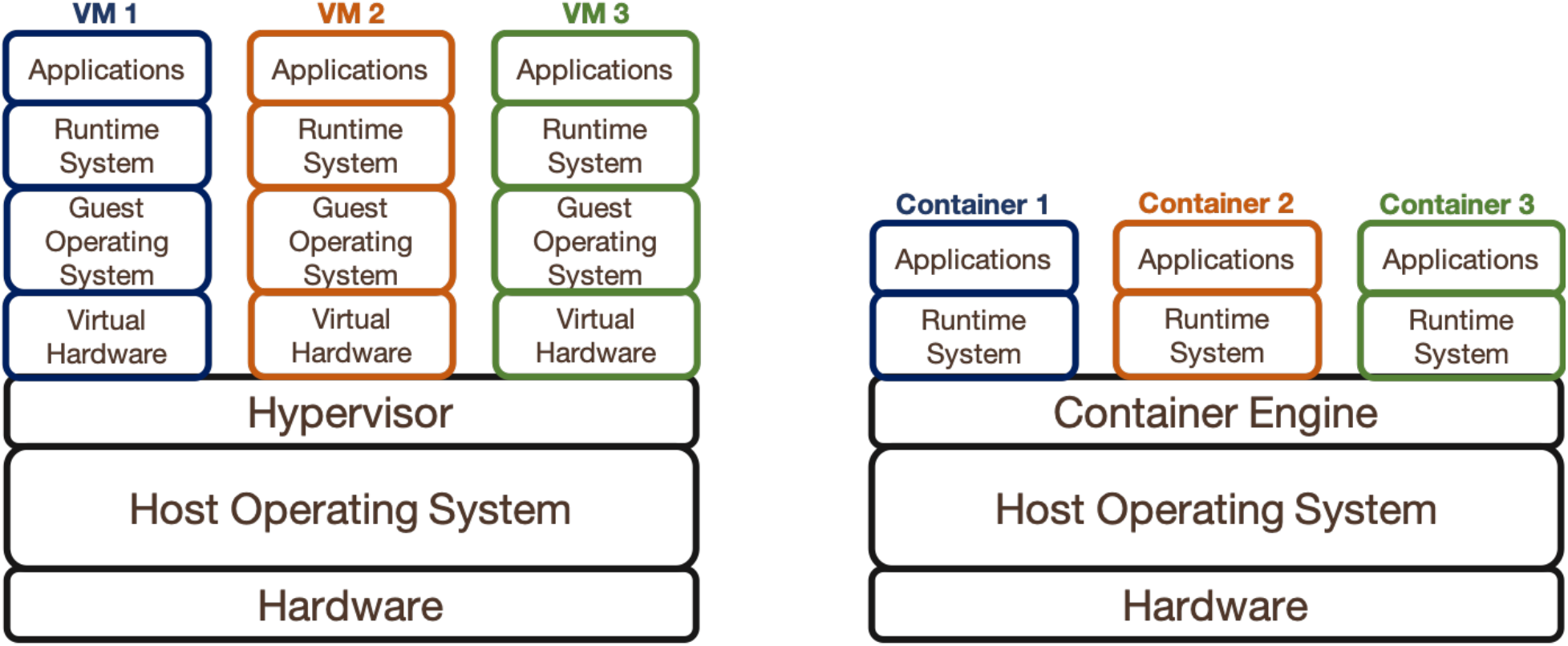
Virtualization vs. Containerization. Virtual Machines (left) require a Hypervisor that creates one or more virtual machines. Each virtual machine comprises virtual hardware that hosts a guest operating system containing the necessary runtime libraries and applications. Containers (right) require a container engine that can carry one or more containers and interfaces these containers with the host OS and hardware. Each container comprises the necessary runtime libraries and the applications.

**Figure 2.**
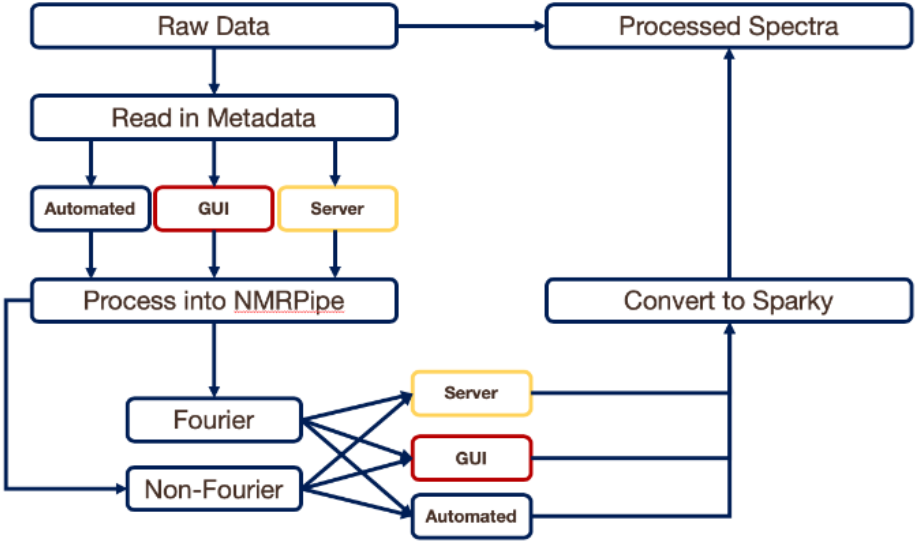
Workflow of NMRdock. The standard workflow of NMRdock allows for the simple processing of raw NMR data through NMRPipe and into Sparky. NMRdock has all of the functionality of both NMRPipe and NMRFAM Sparky and can process both Fourier and NUS data. Spectra can be further analyzed in NMRFAM Sparky.

Several container engines exist, including Rkt, Singularity, and Docker. Unlike with the various implementations of virtual machines, each of these engines has been designed for different purposes and often supports only a single operating system: specifically, Rkt and Singularity both run exclusively on Linux. Fortunately, while Docker runs natively on Linux alone, it has been adapted to run on both Microsoft Windows and Mac OSX, with installers made easily available to end-users (https://docs.docker.com/). Therefore, we utilized the Docker Engine as the base for NMRdock.

## Building NMRdock

The current NMRdock image includes NMRPipe^*25*^ and NMR-FAM Sparky^*35*^ for processing and viewing multidimensional NMR data. All necessary parties have consented to use. NMRPipe is available witout charge from NIST and NIH, while NMR-FAM Sparky is accessible under the GNU General Public License. Within the NMRdock image, NMRPipe and Sparky are both stored in the home directory (/home/ubuntu/). The home directory also includes a Data directory (/home/ubuntu/data) for use as a volume for the container to access data on the host computer. When the NMRdock image is run, the directory that contains the data of interest should be mounted to the image as the data directory in order to allow access. NMRdock can be run in either C-shell or Bash; however, we have optimized the container to run in Bash. To accommodate both shells, we have included a series of wrappers that allow the C-shell (necessary for NMRPipe) to be accessed within Bash. The build files of the current NMRdock image are accessible on GitHub (https://github.com/compbiocore/nmrdock/). We encourage the opening of issues on GitHub for any bugs or requested features.

## Utilizing NMRdock

The general workflow for NMRdock (Figure 2) uses NMRPipe and NMRFAM Sparky to process raw NMR data and to import the processed data into Sparky for analysis. Additional features are being built to expand the base system. NMRdock requires two dependencies: the Docker Engine and an X Window System. The OS-specific Docker Engine (https://docs.docker.com/) must be installed in order to interface the container image with the host operating system. Both NMRPipe and Sparky rely on X Windows for displaying data, and in order to use the GUIs, an XServer is required for your operating system. Furthermore, Mac and Windows operating systems both require a tool for TCP port forwarding to pass data between the container and the host; any such utility can be used for that purpose. On MacOSX, the Socat port forwarder can be installed using Homebrew or MacPorts, and XQuartz is the most widely used implementation of XServer for OSX. We suggest VcXsrv or Xming on Windows, while most distributions of Linux natively include a port forwarder and an X Window system.

There is a quick installation guide at https://compbiocore.github.io/nmrdock/ that will aid in installing Docker and the XServer for your operating system. Detailed instructions for installation of NMRdock are also available. Once NMRdock is installed, it can be accessed from a terminal window or through the executable. It should be noted that changes made within the NMRdock image are not persistent, so all changes (*i.e.* processed and analyzed data) should be stored on the host computer before exiting NMRdock.

## Challenges

One of the major challenges of any software package is longevity. As modern computation evolves, libraries necessary for running older software are either updated or removed. An example that may impact the NMR community is the planned removal of 32-bit support from Mac OSX in Q4 2019 and from Microsoft Windows in 2019 or 2020. NMR software designed and compiled for 32-bit processing will likely experience issues with this transition as computer architectures move exclusively to 64-bit processing. For example, NMRDraw, contained within the NMRPipe package, will no longer be supported natively on Mac OSX. Although we are uncertain of the future changes being made to Mac OSX and Microsoft Windows, the NMRdock container is expected to allow continued use of the 32-bit NMRDraw until it can be replaced by a 64-bit application. The Docker-based NMRdock is stable and continuously supported, where any updates to the base Docker engine will automatically trigger a rebuild of the NMRdock image. NMRdock is also maintained on GitHub, so issues that arise and features to be considered for future builds can be opened as issues directly within GitHub for the developers.

## Educational Tools and Ongoing Development

In addition to its utility for NMR data processing and analysis by intermediate-to-end users through GUIs or the command line, we aim to make NMRdock a widely accessible educational tool for bio-NMR data analysis, where little-to-no up-front computational and coding expertise is required to generate useful results. The NMRdock module may be of particular interest to laboratories that are “light” or “occasional” users of NMR that may not be equipped to install software written in multiple languages or seek out custom processing scripts.

We envision two directions for ongoing work on this module. First, we will work with other developers to increase the library of containers for current and future bio-NMR software. NMRdock is designed to be a modular system, with the base platform described here being supplemented by continued additions of other NMR software packages (Figure 3). This allows the user to easily install and control the necessary NMR packages for their workflow (Figure 4). Second, we will develop and implement automated pipelines for NMR data processing, requiring only the input of raw data files and execution of a single command to generate a complete NMR spectrum. This will allow the creation of server-based NMR data processing suites.

**Figure 3.**
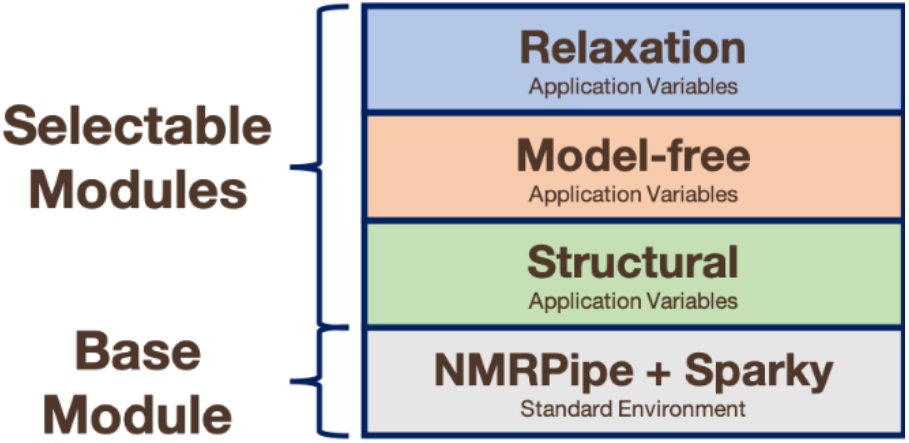
Modularity of NMRdock Containers. NMRdock can be expanded with additional modules. The base NMRdock module presented here can then interface with future modules to aid in the analysis of relaxation, model-free, and structural data.

**Figure 4.**
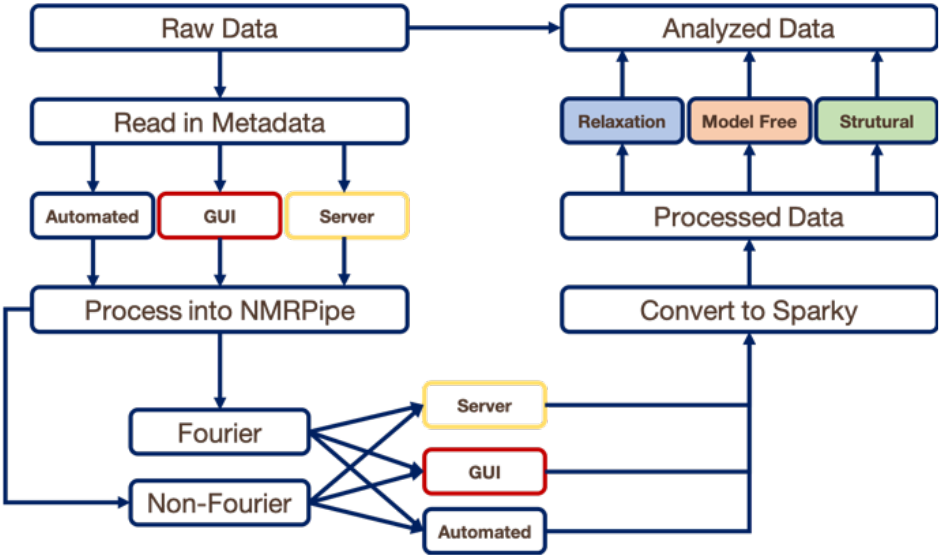
Modular Workflow of NMRdock. Advanced workflows of NMRdock include the current build in addition to future modules that can analyze processed data extracting relaxation, model-free, and structural parameters.

## Conclusions

In this work, we have described a containerization method for the development and maintenance of bio-NMR data processing and analysis software. Using the Docker Engine framework, we have designed NMRdock, containing NMRPipe^*25*^ and NMRFAM Sparky,^*35*^ for the processing of raw NMR data into spectra and the analysis of those spectra. By introducing this tool, we aim to make NMR more accessible for new users, create a long-term solution for aging software, and design an accessible educational tool.

## Acknowledgments

This work was supported by start-up funds from Brown University and funds from the COBRE Center for Computational Biology of Human Disease (P20GM109035) to GPL.

